# Rapid recalibration of temporal order judgements: Response bias accounts for contradictory results

**DOI:** 10.1101/576249

**Authors:** Brendan Keane, Nicholas S. Bland, Natasha Matthews, Timothy J Carroll, Guy Wallis

## Abstract

Recent findings indicate that timing perception is systematically changed after only a single presentation of temporal asynchrony. This effect is known as rapid recalibration. In the synchrony judgement task, similar timing relationships in consecutive trials seem more synchronous (*positive* rapid recalibration; Van der Burg et al., 2013, 2015). Interestingly, the direction of this effect is reversed for temporal order judgements (*negative* rapid recalibration; Roseboom, 2019). We aimed to determine whether *negative* rapid recalibration of temporal order judgements (TOJs) reflects genuine rapid temporal recalibration, or a choice-repetition bias unrelated to timing perception. In our first experiment we found no evidence of rapid recalibration of TOJs, but *positive* rapid recalibration of associated confidence. This suggests that timing perception had rapidly recalibrated, but that this was undetectable in TOJs, plausibly because the *positive* recalibration effect was obfuscated by a large *negative* bias effect. In our second experiment, we dissociated participants’ previous TOJ from the most recently presented timing relationship, mitigating the choice-repetition bias effect, and found evidence of *positive* rapid recalibration of TOJs. We therefore conclude that timing perception is rapidly recalibrated *positively* for both synchrony and temporal order judgements. It remains unclear whether rapid recalibration occurs at the level of sensory processing, leading to similar effects in all subsequent judgements, or reflects a generalised decision-making strategy.

## Introduction

It is well-established that our perception of the relative timing of simple physical events is malleable. For instance, repeated exposure to temporal asynchrony between physical signals (e.g., an audio-track lagging behind the picture at a cinema) leads to a shift in perception such that the events seem to occur more synchronously (e.g., Fujisaki et al., 2004; Vroomen et al., 2004). This effect is known as temporal recalibration, and can be induced experimentally by presenting participants with prolonged periods of asynchronous audio-visual signals. Interestingly, recent work has found that this effect does not require extensive periods of adaptation to manifest, instead, a single presentation is sufficient to influence subsequent timing perception (*rapid* recalibration). However, while temporal recalibration following prolonged exposure to asynchrony is consistently *positive* (timing relationships similar to the adapted asynchrony seem more synchronous), rapid recalibration effects appear to differ based on how participants report their perception of relative timing (Roseboom, 2019).

Van der Burg et al. (2013) found that participants’ judgements of the synchrony of temporally offset audio-visual events is biased toward the temporal relationship presented on the previous trial (for a recent replication, see Figure 1A). That is, participants were more likely to report that an auditory and visual stimulus had occurred simultaneously if they had been presented at a similar timing relationship on the previous trial. Interestingly, this effect was of a similar magnitude to that found following prolonged adaptation (e.g., Vroomen et al., 2004). Van der Burg et al. posited that this effect might reflect a rapid sensory adaptation process. However, the interpretation of these results has been recently challenged (e.g., Roseboom, 2019; Simon et al., 2017).

**Figure 1.**
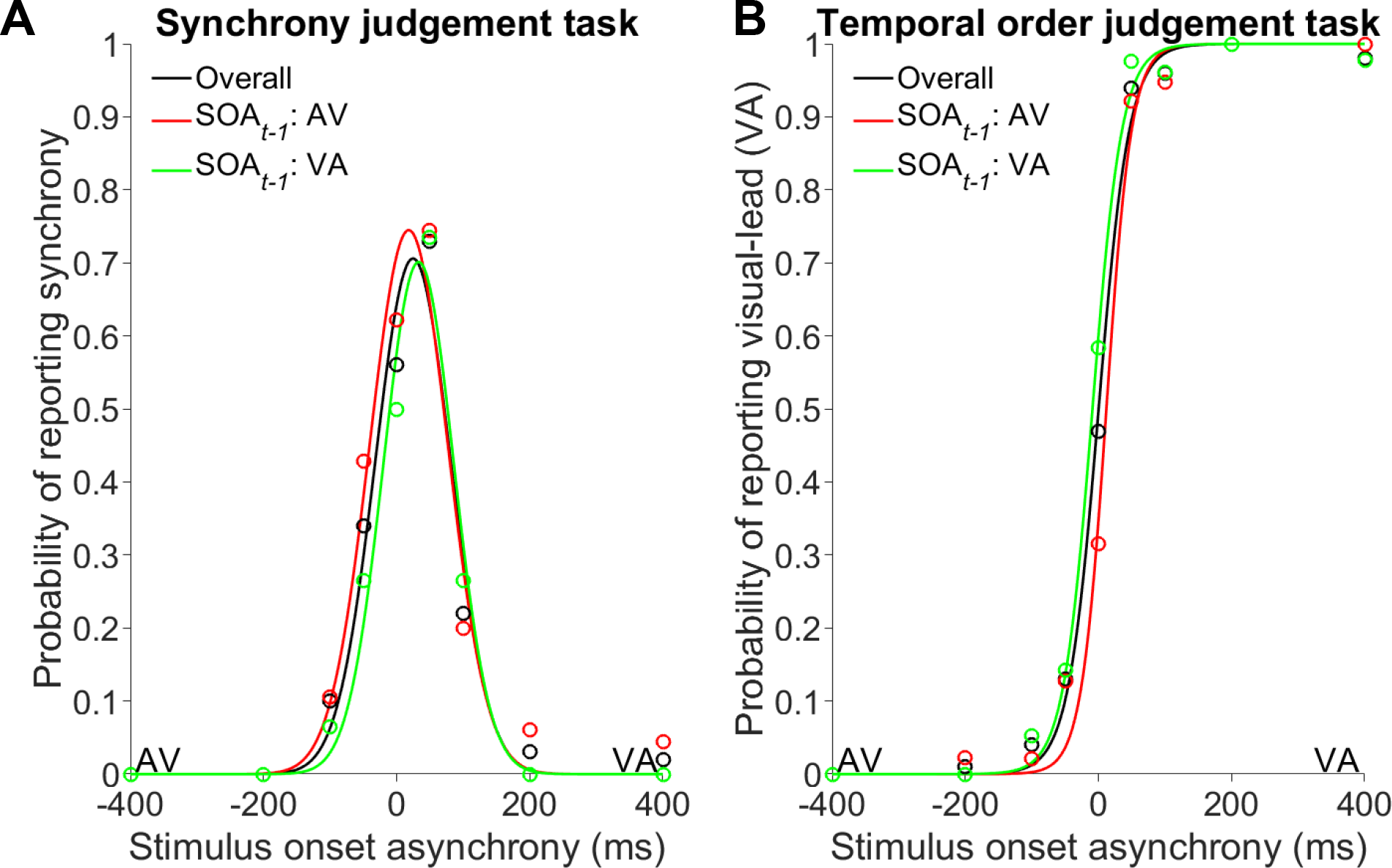
Rapid recalibration of synchrony and temporal order judgements. **(A)** *Positive* rapid recalibration of synchrony judgements, recreated from data provided by Roseboom (2019). Note that this result replicates those of Van der Burg et al (2013, 2015) and Simon et al. (2017). **(B)** *Negative* rapid recalibration of temporal order judgements, recreated from data made available by Roseboom (2019). Three data series are plotted: **black)** overall response distributions, regardless of the stimulus order in the preceding trial; **red)** judgements from trials following an audio-lead (SOA_*t−1*_: AV); **green)** judgements from trials following a visual-lead (SOA_*t−1*_: VA). In both cases, rapid recalibration is measured as a lateral shift of the central tendency of fitted response distributions. The centre of these distributions is generally taken as an estimate of the timing relationship for which the participant is most likely to report perceiving synchrony.

Roseboom (2019) hypothesised that if rapid recalibration reflects adaptation of sensory processing, it ought to occur in other timing tasks, which (presumably) rely on the same sensory processing mechanisms. To test this hypothesis, he had participants complete a synchrony judgement task (SJ; as in Van der Burg et al., 2013), a temporal order judgement (TOJ) task, and a temporal magnitude judgement task. In the SJ task, Roseboom replicated the effects of Van der Burg and colleagues. Participants were more likely to report synchrony when presented with a timing relationship similar to that presented on the previous trial (a *positive* rapid recalibration effect; Figure 1A). However, participants displayed the opposite bias in both the TOJ task (see Figure 1B) and temporal magnitude judgement task (not shown here). That is, participants were *less* likely to perceive synchrony when presented with similar timing relationships in consecutive trials (a *negative* rapid recalibration effect). Roseboom concluded that a purely sensory account of rapid recalibration cannot explain these discrepant effects, assuming that some sensory process generates a timing estimate which is then evaluated by subsequent decision-making processes. Instead, he proposed that rapid recalibration reflects changes in decisional processes, and these changes manifest differently for synchrony and temporal order judgement tasks.

The discrepancy in the direction of effects reported in TOJ and SJ tasks is difficult to reconcile with a purely sensory account of rapid recalibration. Assuming rapid recalibration induces changes in timing perception consistent with those induced by prolonged exposure to asynchrony, both tasks should reveal *positive* rapid recalibration effects. However, since rapid recalibration effects depend on the previous trial, any serial dependency in the participants’ responses may impact the observed recalibration effect. Unfortunately, the way in which rapid recalibration is determined for order judgement tasks (TOJ, and temporal magnitude judgements) means the effect is vulnerable to contamination by choice-repetition biases (Pape et al., 2017). Critically, this problem is not present in the SJ task.

In general, participants are reasonably sensitive and accurate when making judgements of stimulus order (e.g., Donohue et al., 2010). If the visual stimulus was leading on the current trial, the participant would likely report this. In the subsequent trial, regardless of the timing of the stimuli, the simple bias to repeat their previous TOJ inclines the participant to report a visual-lead stimulus-onset asynchrony (SOA) again. This uniformly increases their probability of reporting a visual-lead for any SOA following a visual-lead. The consequence, then, is that the area under their discriminant function increases, shifting their point of subjective synchrony (PSS) toward audio-lead SOAs (see simulated response distributions in Figure 2B).

**Figure 2.**
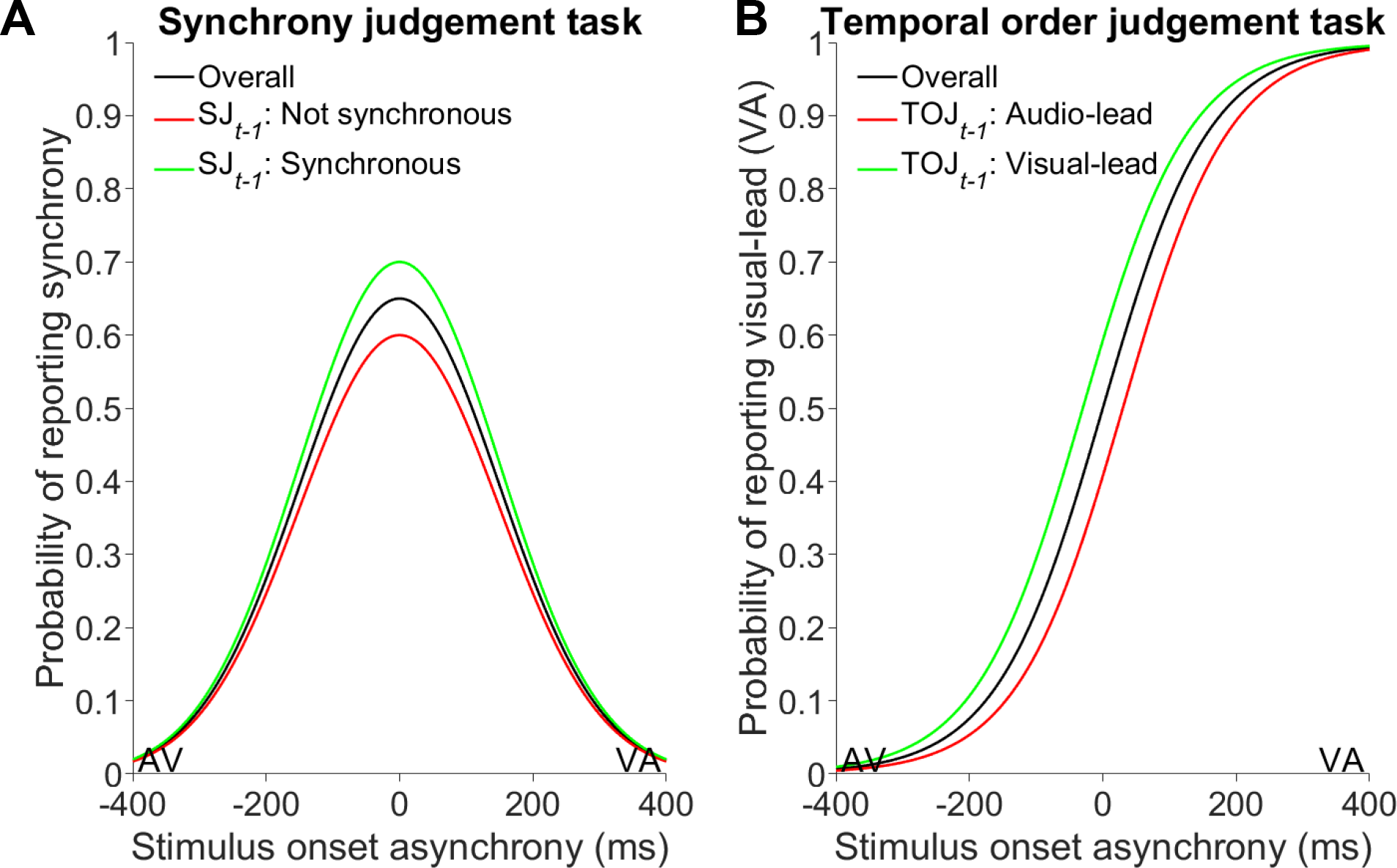
*Simulated choice-repetition bias effects in synchrony and temporal order judgement tasks*. Plotting conventions are the same as those used in Figure 1, except the judgements are split by the preceding judgement (i.e., SJ or TOJ on trial *t−1*) rather than the preceding timing relationship. **(A)** Choice-repetition bias in synchrony judgements produces a relative vertical shift of response distributions, with no change in lateral position of the distribution (remembering that a horizontal shift would indicate recalibration). **(B)** Choice-repetition bias in temporal order judgements produces a lateral shift of response distributions, mimicking the *negative* rapid recalibration effect reported by Roseboom (2019).

Similarly, if the previous trial was an audio-lead, the participant probably chose to report an audio-lead, and a choice-repetition bias would slightly increase the probability the participant would do so again in the subsequent trial. At the block level, this would produce a PSS for trials following audio-lead SOAs closer to visual-lead SOAs. Working in concert, a bias to repeat choices would produce an effect in TOJ tasks that mimics the *negative* recalibration effect (Figure 2B) reported by Roseboom (2019). Therefore, we cannot be sure whether the effect reported by Roseboom is due to the timing of stimuli presented in the previous trial, or a simple bias to repeat TOJs. In essence, the previous choice and the previous SOA are confounded. Note that this choice-repetition bias would also produce changes in SJ distributions, but only in terms of amplitude, with no change in PSS (in this case taken as the mean of the distribution; see Figure 2A). This difference in how a choice-repetition bias manifests for these two tasks may explain the discrepancy in reported rapid recalibration effects.

Here we aimed to resolve the key issue of whether *negative* rapid recalibration of TOJs reflects a choice-repetition bias, unrelated to timing perception. We first show (*in silico*) that *negative* rapid recalibration effects may manifest during TOJ (but not SJ) tasks as a result of a choice-repetition bias. We then show that this bias is present in Roseboom’s data, and predicts his rapid recalibration effects in temporal order and magnitude judgement tasks. In Experiment One we used confidence as a secondary probe of timing perception. We failed to replicate rapid recalibration of TOJs, but found evidence of *positive* rapid recalibration in associated confidence judgements. In Experiment Two, we aimed to dissociate the preceding SOA from participants’ preceding TOJ, thereby mitigating the effect of a choice-repetition bias on TOJs. Using this paradigm, we found evidence of *positive* rapid recalibration of TOJs. The results of these studies indicate that timing perception rapidly recalibrates consistently across timing tasks.

### Choice-repetition bias analyses

Roseboom (2019) reported a *negative* rapid recalibration effect in temporal order and temporal magnitude judgement tasks, while replicating the previously-reported *positive* rapid recalibration effect in synchrony judgements (Van der Burg et al., 2013, 2015). A simple choice-repetition bias (Pape et al., 2017) could account for these discrepant results, considering the shape of the associated response distributions (see Figure 2). Van der Burg et al. (2013) analysed participants’ SJs as a function of their preceding judgement, and found no effect on PSS estimates, consistent with the idealised curves presented in Figure 2. These control analyses were not reported by Roseboom (2019). However, Roseboom has made the raw data from his experiments available online.

We analysed Roseboom’s (2019) data and found striking biases to repeat choices on consecutive trials in both the SJ and TOJ tasks (and the magnitude judgement task; not shown here). In the SJ task, we compared participants’ judgements following trials where they reported synchrony against those following trials where they reported asynchrony (Figure 3A). Similarly, in the TOJ task, we compared participants’ responses on trials following an audio-lead (AV) report versus following a visual-lead (VA) report (see Figure 3B). The choice-repetition bias manifested as a change in the amplitude of the SJ distribution (as idealised in Figure 2A), with no systematic change in PSS. However, this bias shifted the PSS in the TOJ task, consistent with *negative* rapid recalibration.

**Figure 3.**
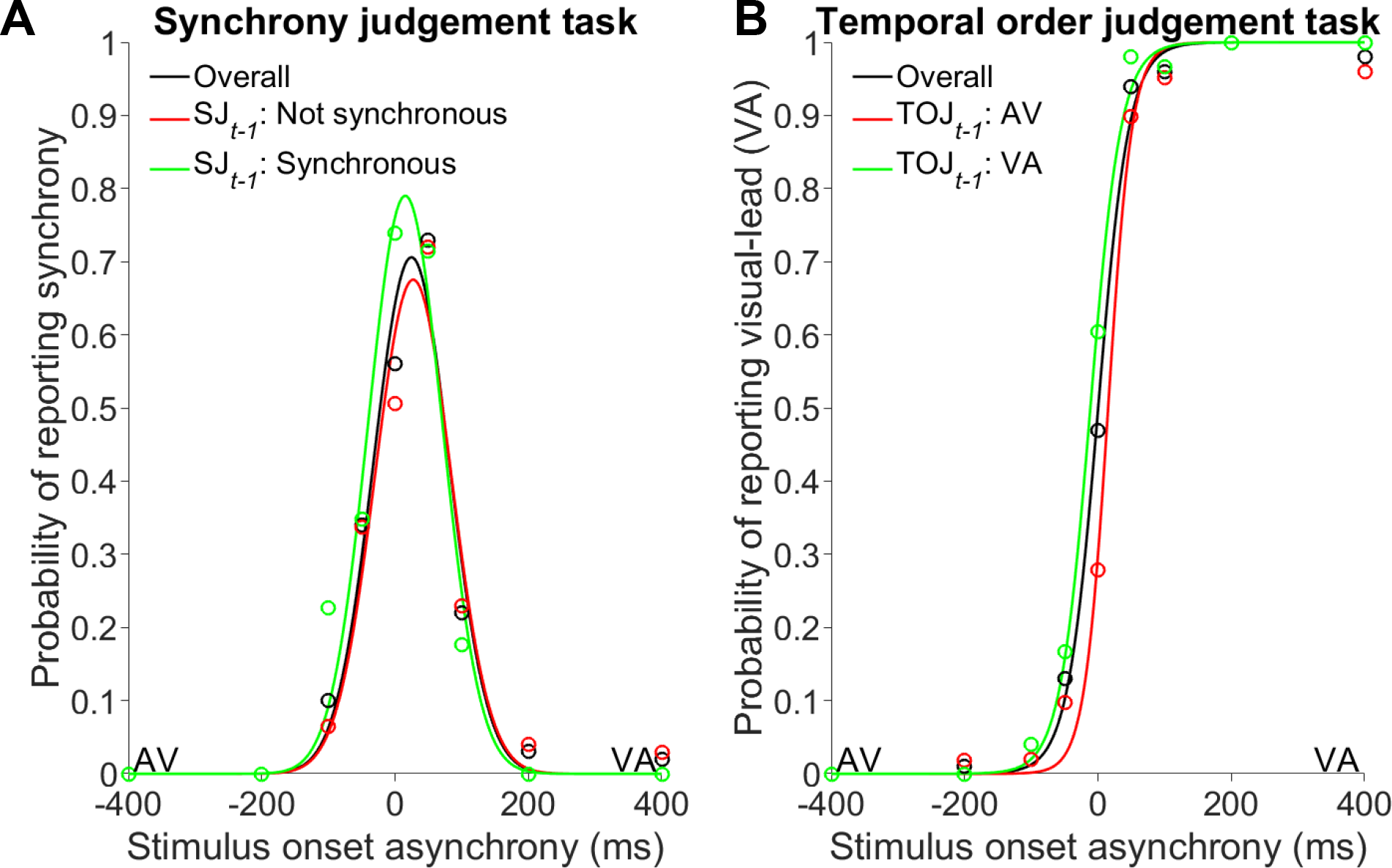
*Choice-repetition biases in synchrony and temporal order judgements*. These images are generated from re-analysis of data collected in Roseboom (2019). Plotting conventions are the same as those used in Figure 2. **(A)** A choice-repetition bias produces a shift only in the amplitude of the synchrony judgement response distribution, but not the mean. In this sense, the synchrony judgement task prevents any simple response bias (choice-repetition or otherwise) from producing false rapid recalibration effects. **(B)** A choice-repetition bias in the temporal order judgement task produces shifts in the mean of the response distribution, unlike the synchrony judgement response distribution. Comparison of distributions reveals a *negative* rapid recalibration effect of temporal order judgements (despite being analysed in terms of preceding judgement, rather than preceding timing relationship). Note that Figures 1B and 3B are not identical, though their similarity is striking.

We then investigated the contribution of choice-repetition bias effects to the apparent rapid recalibration effects. For each participant we calculated their rapid recalibration effect by computing the difference in their PSS (see Methods, Equation 1) for TOJs following AV trials versus those following VA trials. We computed their choice-repetition bias effect using the same logic, but with trials split by the preceding TOJ, rather than preceding stimulus-order. In both the temporal order and magnitude judgement tasks, paired *t*-tests indicate that the choice-repetition bias effect (*M* = −99.00ms, *M* = −74.26ms, respectively) is significantly larger than the rapid recalibration effect (*M_TOJ_*= −34.04, *t*_19_ = −3.80, *p* = .001; *M_Mag_* = −41.46ms, *t*_17_ = −3.10, *p* = .006). Further, the magnitude of these two effects was positively correlated in the TOJ (*r_18_* = 0.81, *p* < .001) and magnitude judgement tasks (*r_16_* = 0.66, *p* = .003), even capturing the *positive* rapid recalibration of participants with choice-alternation biases.

We then attempted to quantify the magnitude of the choice-repetition bias required to elicit a false rapid recalibration effect. We simulated observers in a TOJ task with varying probabilities of simply repeating their preceding TOJ. On each trial, a simulated observer perceived an SOA drawn from a random normal distribution with a mean of the actual SOA and variance of 80ms, and compared this to a fixed criterion at 0ms to make a temporal order judgement. On some proportion of trials, the simulated observer simply repeated the previous TOJ, regardless of SOA. We simulated 20 of these observers completing an experiment using the same SOAs as Roseboom (2019), with the same number of trials, split into the same number of testing blocks. This experiment was simulated 1,000 times per repetition rate. We found that the proportion of trials required to produce significant rapid recalibration effects was alarmingly low (see Figure 4).

**Figure 4.**
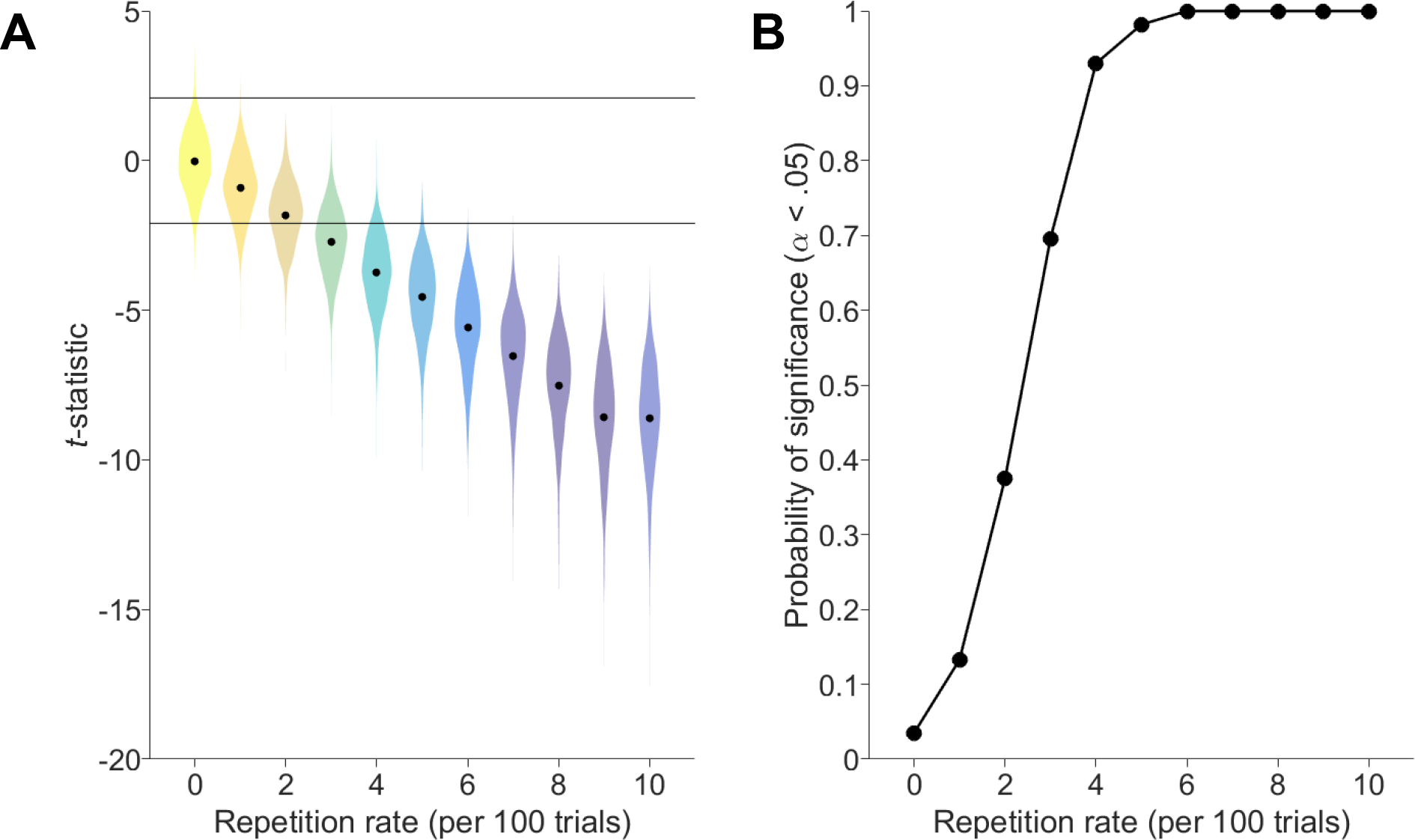
Results of simulated experiments. We simulated the effect of choice-repetition rates ranging from 0 to 10 repetitions per 100 trials, in experiments with a sample size of 20. Note that each participant in these simulated experiments had the same repetition-rate and perceptual sensitivity, but a random trial order. **(A)** We conducted 1,000 virtual experiments for each repetition-rate, and tested for rapid recalibration (see Methods, Equations 1 & 2). Black horizontal bars indicate the *t*-statistic corresponding to significance (α < .05) in a paired-samples *t*-test with *df* = 19. **(B)** The proportion of *p*-values from these *t*-tests that exceed the significance criterion. That is, the probability of finding a significant rapid recalibration effect as a function of repetition-rate. Note that the simulated observer was coded to make completely independent responses across trials (i.e., no form of ‘rapid recalibration’ was built into the simulated observer), except for the variable proportion in which they simply repeated their preceding TOJ regardless of current-trial SOA.

We found that the probability of finding a significant ‘rapid recalibration’ effect exceeded 90% with a choice-repetition rate of only 4%. That is, if participants repeat their preceding TOJ on only 4 out of every 100 trials, perhaps due to lapses in concentration, there is a ~93% chance of finding evidence of a *negative* rapid recalibration effect. This is alarming, and plausible. Pape et al. (2017) found that participants were strongly biased to repeat their orientation judgements in consecutive trials, even with intervening button-presses and a random response-mapping on each trial. Verstynen and Sabes (2011) asked participants to reach to targets incrementally rotating around a circular track. Despite the target location being perfectly predictable, participants were biased to reach toward the location where the target had just been. Marinovic et al., (2017) reported a similar effect, where participants were consistently biased toward an irrelevant location to which they had previously moved, even when given explicit instruction that the location was irrelevant. The bias effect remained even when participants were provided a long time (~ 1s) to prepare each movement after target presentation. These results suggest that choices, and the motor behaviours that enact them, are reliably biased toward preceding instances, despite explicit instructions, predictable patterns, and ample preparation time.

We took a very simple approach to quantifying the repetition rate in Roseboom’s (2019) data. In the temporal order and magnitude judgement tasks, we computed the probability of each response being the same as the response made on the preceding trial (based only on stimulus-order for the magnitude judgement task). We then subtracted the probability of the correct response repeating in consecutive trials, to estimate the repetition bias over and above the repetition built into the actual stimulus order. In temporal order and magnitude judgement tasks, we found that Roseboom’s participants repeated their responses on ~56% of trials, while the correct response (dictated by the stimulus order) was repeated only ~50% of the time (as expected). According to our simulations, a 6% choice-repetition rate is extremely likely (>99%) to produce false evidence of *negative* rapid recalibration. In fact, of our 1,000 simulations of a repetition rate of 6%, every single sample returned a significant *negative* rapid recalibration effect.

In our first experiment, we aimed to replicate the reported *negative* rapid recalibration of TOJs, while also measuring participants’ confidence in their TOJs. This design allows us to exploit the shape of the confidence distribution, which ought to be immune to a choice-repetition bias (as in SJ tasks). If timing perception rapidly recalibrates *positively*, similar timing relationships in consecutive trials should seem more synchronous, or less ordered, and so be associated with lower confidence. In essence, confidence in timing judgements should rapidly recalibrate in line with rapid recalibration of timing perception itself.

### Experiment One methods

Fifty adults (21 male, 29 female; ages ranged 18–33 years, *M* = 21.83) were paid to participate in a single one-hour testing session involving an audio-visual TOJ task. Participants gave written informed consent before testing started, and were remunerated $20/hour for their time. This study was approved by the ethics committee of the School of Human Movement and Nutrition Sciences at The University of Queensland. Participants reported having normal or corrected-to-normal hearing and vision, and were naïve to the purposes of the experiment. Two participants were removed from analyses because their sensitivity to temporal order was extremely poor (just-noticeable difference greater than the sampling range, indicating they could not reliably discern the temporal order of even the most asynchronous stimuli, offset by 300 ms).

Note that the sample of participants analysed here is aggregated from two iterations of this experiment. The first sample consisted of 20 participants, completing 600 trials of a TOJ task with a confidence judgement. We decided to re-run this study with another (independent) sample, with 50% more participants (*N* = 30), and 20% more trials (720). These participants also completed a short familiarisation block of trials before starting the experiment, with no feedback, to give them the opportunity to ask specific questions about the task. We believe these samples are appropriately matched in terms of experimental procedures and demographic properties. Analysis of each sample separately yields results consistent with the aggregate sample, and so for simplicity we report the pooled samples.

Participants sat in a dimly lit room with their chin placed in a chin rest 57 cm from the computer monitor. A small red LED (0.5 cm diameter; 0.5 degrees of visual angle, DVA) was attached to the centre of the monitor, and speakers were hidden behind the monitor. On each trial the red LED was illuminated for 20 ms, and the speakers played a square-wave tone of 440 Hz for 20 ms. The auditory stimulus was onset at one of six SOAs relative to the visual stimulus (±300, ±200, and ±100 ms; i.e., never synchronous). Each SOA was sampled 100 times in pseudorandom order for the first 20 participants, and 120 times each for the remaining 30 participants. Trial structure is schematised in Figure 5. Audio and visual stimuli were generated by an independent microcontroller. Stimulus presentation timing was validated prior to data collection using an oscilloscope, and also measured during testing using another independent microcontroller. Online measurement allowed for rejection of trials in which stimulus presentation timing deviated from intended timing. Note that no trials were rejected on this basis. Stimulus presentation timing did not deviate from intended timing by more than 0.24 ms on any trial for any participant (*M* = –6 μs, *SD* = 86 μs).

**Figure 5.**
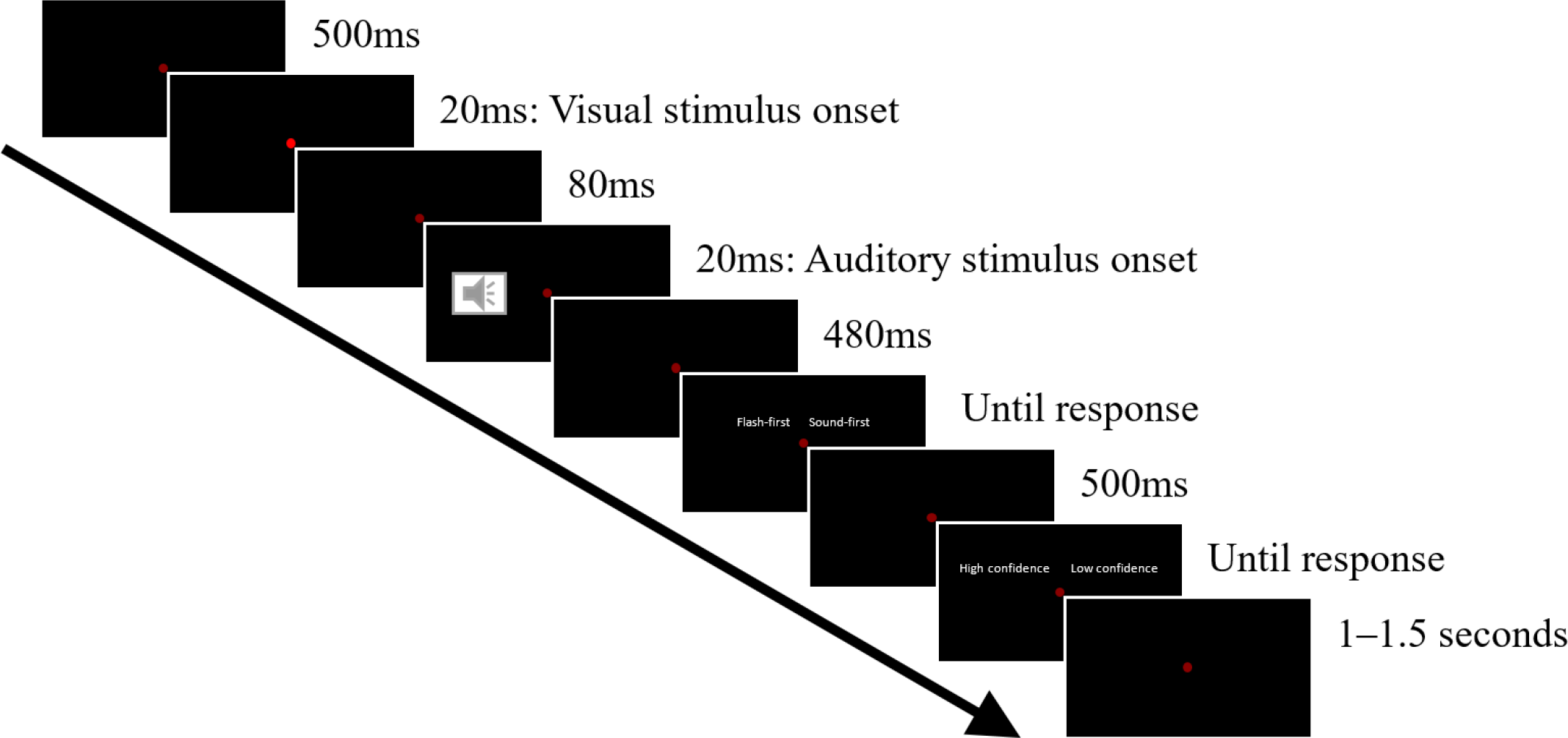
Schematic of the trial structure, with an SOA of 100ms. Each trial was preceded by a period of silence and a blank screen lasting between 1 and 1.5 seconds. The trial period then started, lasting a total of 1.1 seconds. The visual stimulus appeared 500 ms into the trial period on every trial, while the auditory stimulus onset at any of the six SOAs relative to the visual stimulus (±300, ±200, and ±100 ms). Note that the ‘loudspeaker’ symbol appearing in the figure is diagrammatic only and no visual cue was provided with audio onset. Note that the red LED was always visible, but only became illuminated (here in the second panel) to indicate the visual signal (i.e., it was otherwise dim).

Following stimulus presentation, participants were asked to report which of the audio and visual stimuli had occurred first. Participants were prompted by the appearance of a text message on screen behind the (now dim) red LED, informing them to press either the left or right mouse button to indicate their choice. Participants were then prompted to report their confidence (either high or low; see Figure 5) in their TOJ. Response mappings for temporal order and confidence judgements were counterbalanced across participants. Participants were given a one minute break every 100 trials (approximately every seven minutes). The first trial, and the trial immediately following each break, were excluded from analyses since no trial preceded them.

### Data Analysis

In order to determine the extent to which each TOJ was influenced by the temporal order of stimuli on the preceding trial, participants’ temporal order and confidence judgements were split into two datasets: trials in which the visual stimulus physically occurred first in the preceding trial (SOA_*t−1*_: VA), and trials where the visual stimulus physically occurred second in the preceding trial (SOA_*t−1*_: AV). We fitted descriptive functions to response distributions in these datasets, and to participants’ overall dataset (regardless of stimulus order on the preceding trial), and compared model parameters. This analytic approach is consistent with that of recent similar work (e.g., Simon et al., 2017; Van der Burg et al., 2013).

We fitted a logistic function to participants’ TOJs from each dataset (see Eq 1). This provided an estimate of each participants’ point of subjective synchrony (PSS), and their sensitivity to temporal order (just-noticeable difference, JND). We calculated the extent to which the participants’ perception of synchrony was influenced by the temporal order of stimuli on the preceding trial by subtracting the two PSS estimates (see Eq 2).

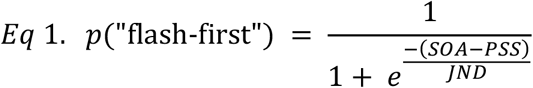

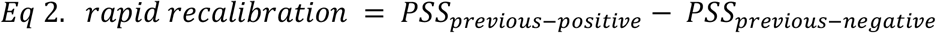

We used a very similar process to quantify the effect of temporal order on the preceding trial on participants’ confidence judgements. We fitted a Gaussian distribution (see Eq 3) to each participant’s probability of reporting low-confidence at each of the six SOAs, for the two datasets. This provided an estimate of the timing relationship for which the participant was least confident (point of least confidence, PLC), and the variance in their low-confidence responses (*σ*). As for order judgements, we estimated the effect of the preceding stimulus-order on confidence judgements by computing the difference in PLCs.

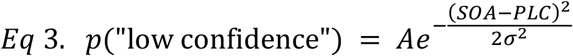

## Experiment One results

### Rapid recalibration analysis

We first calculated participants’ overall bias and sensitivity (PSS and JND, respectively) to audio-visual temporal order. On average, participants were accurate (mean PSS = 20.99 ms, SD = 102.08 ms) and precise (mean JND = 116.97 ms, SD = 73.63 ms). Paired *t*-tests indicated that the temporal order of stimuli on the preceding trial had no systematic effect on participants’ perception of synchrony (*t_47_* = −1.81, *p* = .088), though this *negative*-going effect approached significance. Similarly, we found no evidence that stimulus-order on the preceding trial influence participants’ sensitivity to temporal order (as predicted; *t*_47_ = −1.47, *p* = .158). Contrary to predictions, and unlike previous reports (e.g., Van der Burg et al., 2013), participants’ sensitivity to temporal order was uncorrelated with the magnitude of their rapid recalibration effect (*r_46_* = −0.05, *p* = .761). Participants’ temporal order and confidence response distributions are presented in Figure 6.

**Figure 6.**
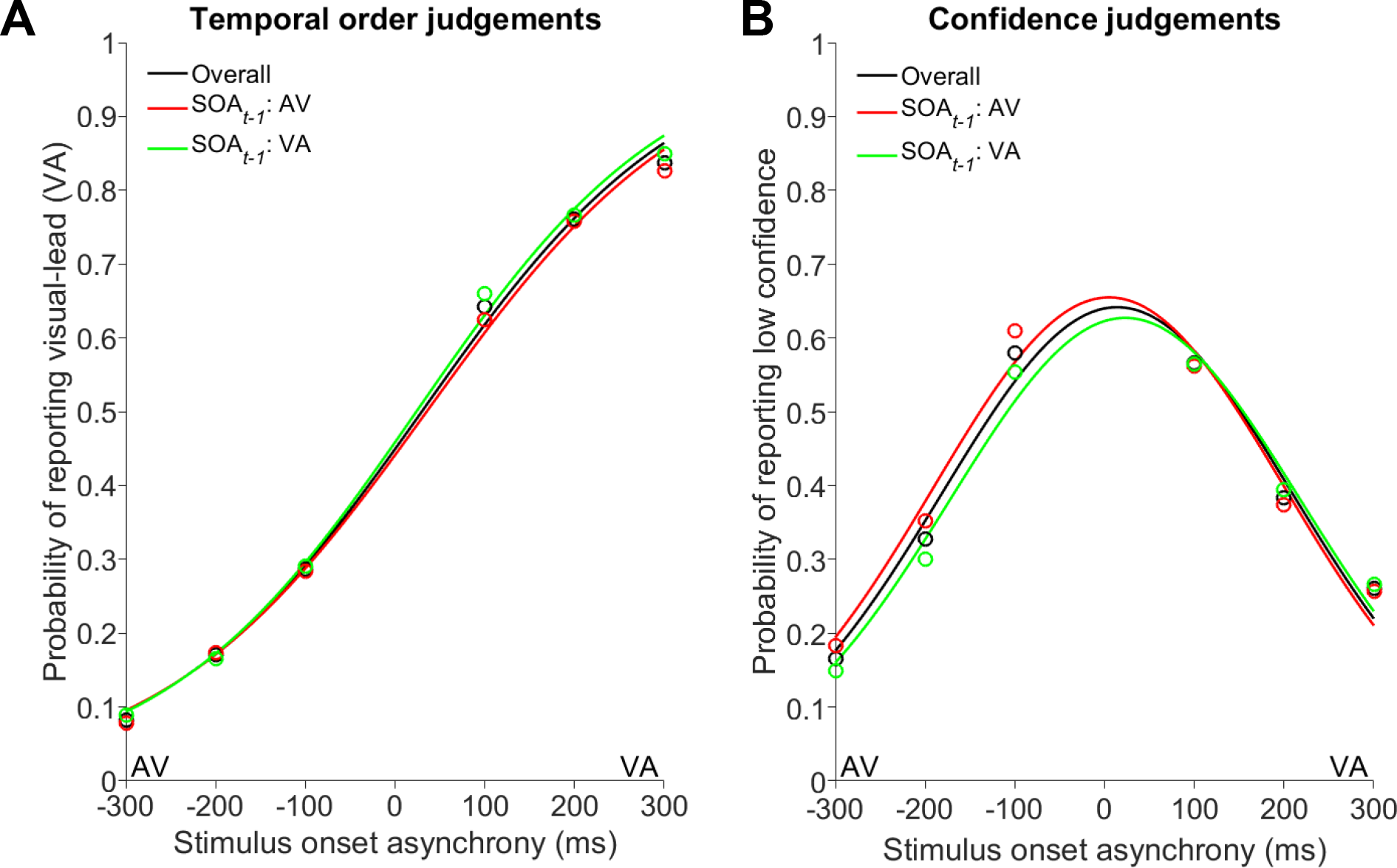
Temporal order and confidence response distributions, collated across participants. Plotting conventions are the same as those used in Figure 1. Note that in all analyses presented here, responses are split into separate datasets based on the physical timing of stimuli in the preceding trial, regardless of the participants’ judgement of stimulus timing on the preceding trial. **(A)** Participants’ temporal order judgements indicate sensitivity to physical timing, as expected. However, there is no evidence of an effect of the order of stimuli on the preceding trial, contrary to predictions. **(B)** Overall, participants were most likely to make TOJs with low confidence when the physical timing offset was near zero, as expected. Participants were more likely to report low confidence for timing relationships similar to that presented in the preceding trial. This is evidenced by a significant difference between the mean of the red and green distributions, corresponding to trials following audio-leads and audio-lags, respectively. This is consistent with a *positive* rapid recalibration of timing perception, with similar SOAs in consecutive trials seeming more synchronous, or less ordered, and therefore judged with lower confidence.

### Confidence analysis

We then fit Gaussian functions (see Eq 3) to participants’ probability of reporting low confidence across each of the six SOAs, independently for each of the two datasets (SOA_*t−1*_: VA, and SOA_*t−1*_: AV). This provided an estimate of the point at which each participant was least confident (PLC), and the variance in their confidence judgements. On average, participants reported least confidence for 18.34 ms visual-leads (SD = 53.26 ms), indicating they were sensitive to the difficulty of the task (mean *σ =* 196.86 ms, SD = 97.19 ms). One participant (in addition to the two removed from TOJ analyses due to poor JNDs) was removed from analyses as the variance in their confidence judgements indicated random responding (*σ* > 1 million).

Paired *t*-tests indicated that the temporal order of stimuli on the preceding trial reliably influenced participants’ PLC (*t_46_* = 2.96, *p* = .009). As predicted, this test indicated no effect of the temporal order of stimuli on the preceding trial on the variance in participants’ confidence judgements (*t_46_* = 0.76, *p* = .460). We found no evidence for a relationship between participants’ recalibration of low confidence and the variance in their reports of low confidence (*r_45_* = 0.10, *p* = .499).

### Choice-repetition bias analysis

We then quantified the effect of recent responses on participants’ TOJs and associated confidence. Figure 7 shows participants’ TOJs (top row) when split by their preceding TOJ (Figure 7A), and their confidence in the preceding TOJ (Figure 7B). Note that their confidence judgement for the preceding TOJ (Figure 7B) corresponds to the button-press immediately preceding the plotted TOJs. Note that two additional participants were removed (as well as those removed from previous analyses) due to poor model fits. Paired *t*-tests on model parameters indicated that participants’ PSS varied as a function of preceding-TOJ (Figure 7A; *t*_44_ = −3.89, *p* = .001), but not preceding confidence (Figure 7B; *t*_44_ = 0.25, *p* = .806). This is consistent with participants’ being biased to repeat their most recent TOJ (and not their most recent button-press). In Roseboom’s (2019) data, we found a strong correlation between the rapid temporal recalibration and response bias effects, in both TOJ and temporal magnitude judgement tasks (*p*s < .05). Similarly, we found that participants’ choice-repetition bias (change in PSS as a function of preceding TOJ) was extremely highly correlated with their rapid recalibration effect (*r*_44_ = 0.82, *p* < .0001).

**Figure 7.**
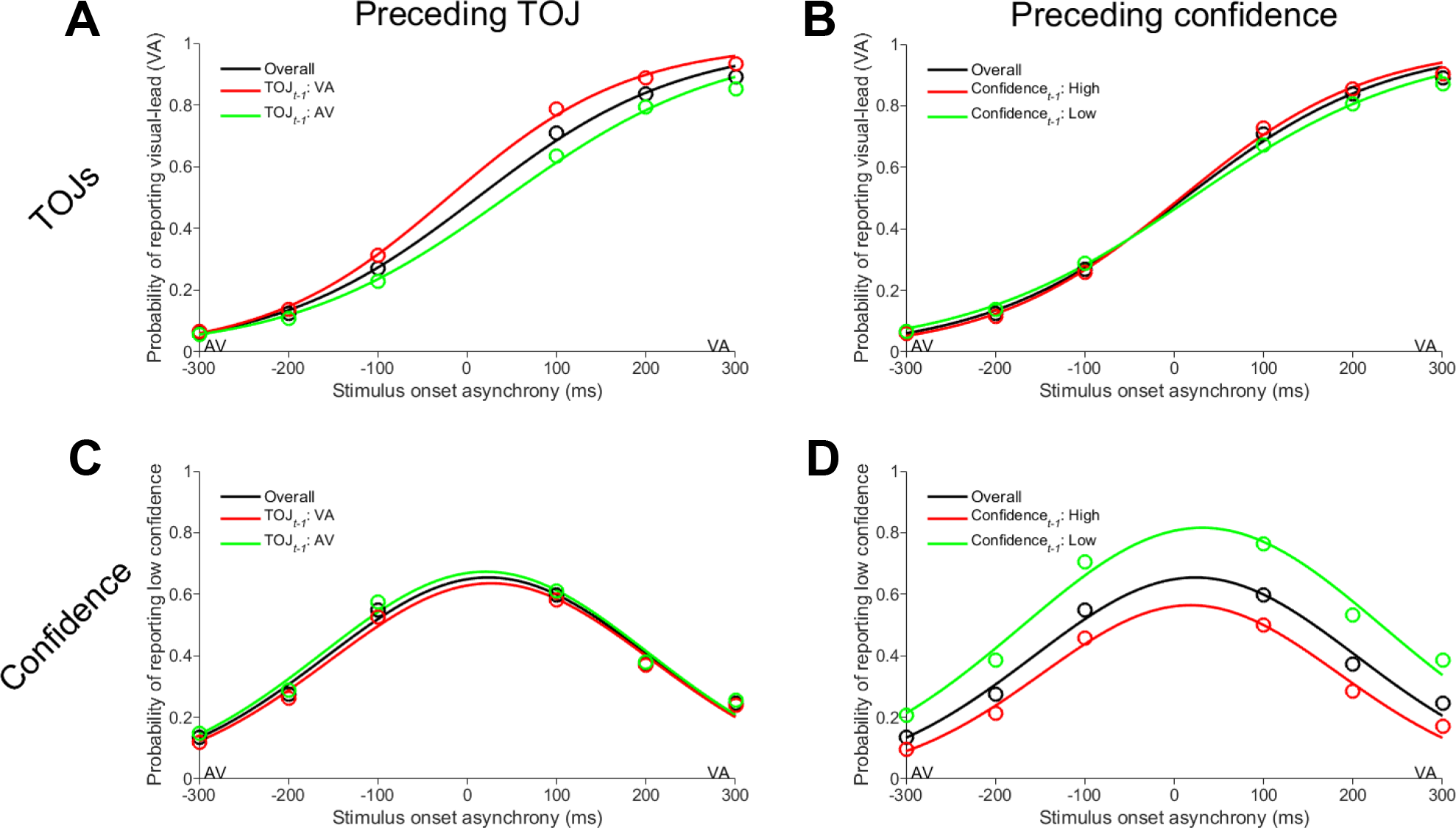
Participants’ judgements as a function of preceding judgements. In order to investigate choice-repetition biases, we split participants’ temporal order and confidence judgements by the preceding temporal order and confidence judgement. **Top row)** Participants’ TOJs as a function of their preceding TOJ and confidence judgement. Note that the preceding confidence judgement is the button-press immediately before these order judgements are made. **(A)** We found an effect of the preceding TOJ on subsequent TOJs. When the participants’ preceding TOJ was a visual-lead (TOJ_*t−1*_: VA), participants were more likely to report a visual-lead again. The same is true for TOJs following an audio-lead judgement. **(B)** However, we found no effect of the preceding confidence report on subsequent TOJs, despite it being the most recent button-press. **Bottom row)** Participants’ confidence judgements as a function of the preceding TOJ and confidence. **(C)** Participants’ previous TOJ did not bias confidence judgements, despite evidence that the preceding SOA influenced confidence (Figure 6B). This suggests that the observed rapid recalibration of low-confidence is not due to a simple response bias. **(D)** Participants’ confidence on the preceding trial, however, did influence their subsequent confidence. Specifically, when participants reported low-confidence on some trial, they were much more likely to do so again on the subsequent trial.

Using the same analytic approach, we then quantified the effect of participants’ preceding responses on their decisional confidence. Figure 7 shows participants’ probability of reporting low confidence (bottom row), as a function of their preceding TOJ and confidence. A similar set of results emerges for confidence judgements; participants’ confidence was unchanged by their preceding TOJ (in terms of amplitude, mean, and standard-deviation; Figure 7C; *p*s > 0.05), but varied reliably as a function of their preceding confidence (Figure 7D). Participants were marginally more likely to report low-confidence if they did so on the previous trial (*t*_44_ = 1.87, *p* = .068), and produce broader low-confidence distributions (*t*_44_ = 2.23, *p* = .031), with no change in PSS (*t*_44_ = 1.29, *p* = .203). Similar to the bias observed for TOJs, this indicates a bias to repeat their preceding confidence judgement.

### Experiment One discussion

We aimed to investigate the effect of stimulus order on subsequent timing judgements, and associated confidence. If temporal order perception rapidly recalibrates positively, in line with synchrony perception (e.g., Van der Burg et al., 2013, 2015), similar SOAs in consecutive trials would appear more synchronous, shifting the TOJ PSS toward the preceding SOA, and lowering participants’ confidence (as they now seem less ordered). Alternatively, if the effect reported by Roseboom (2019) reflects genuine rapid recalibration of temporal order perception, participants’ TOJ PSS should shift away from the preceding SOA, and confidence ought to be increased for judgements of similar SOAs in consecutive trials (as they seem less synchronous, or *more* ordered). We found no evidence of rapid recalibration of temporal order perception, consistent with neither hypothesis, despite using a much larger sample size than most, and comparable trial numbers.

However, we found evidence that confidence judgements varied from trial-to-trial, consistent with the effect predicted by *positive* rapid recalibration (Simon et al., 2017; Van der Burg et al., 2013, 2015). Participants were generally less confident in their TOJs if similar SOAs occurred in consecutive trials, plausibly because they seemed more synchronous. This suggests that temporal order perception *positively* rapidly recalibrated; why then was this effect not detected in the TOJs? We propose that *positive* rapid recalibration of temporal order perception was obfuscated in TOJs by a *negative* response bias. Specifically, a bias to repeat TOJs in consecutive trials. We found clear evidence of this bias in terms of both temporal order and confidence judgements (see Figure 7), and found that the magnitude of this bias in TOJs was positively correlated with individuals’ rapid recalibration effect. Even a small bias of this sort could produce trial-order effects in a TOJ task (see Figure 4) that are erroneously interpreted as rapid recalibration, but have no basis in timing perception. It is therefore perhaps surprising that we did not find a *negative* recalibration-like effect here due to the choice-repetition bias alone (as our simulations would indicate is extremely likely). We therefore reason that the null result reported here is due to the two opposing effects (i.e., *positive* rapid recalibration of temporal order perception and *negative* choice-repetition bias) nullifying one another.

In our second experiment we aimed to dissociate participants’ preceding TOJ from the preceding timing relationship. In doing so, we disrupted the relationship between the previous judgement and the previous SOA present in Experiment One, and Roseboom’s (2019) TOJ and temporal magnitude judgement tasks. Consistent with the results of Van der Burg et al. (2013, 2015), Simon et al. (2017), and our own confidence results here, we predicted that mitigating this choice-repetition bias effect would allow us to detect *positive* rapid recalibration of TOJs.

## Experiment Two methods

We recruited 30 participants to complete an orientation judgement task interleaved with an audio-visual TOJ task. Participants gave written informed consent, and were remunerated $20/hour for their time. This study was approved by the ethics committee of the School of Human Movement and Nutrition Sciences at the University of Queensland. Participants reported having normal or corrected-to-normal hearing and vision, and were naïve to the purposes of the experiment.

On every trial, participants were asked to fixate a red dot, subtending 0.25 DVA, in the centre of the screen. They were then presented with an oriented Gabor patch (12.5 DVA diameter) in a Gaussian aperture (standard deviation of 1.56 DVA) and an auditory tone (440 Hz). Stimuli were 10 ms in duration (corresponding to one frame at 100 Hz). The Gabor was oriented at one of six pre-defined orientations relative to vertical (±4, ±2, and ±1 degrees). Presentation of the audio and visual stimuli was asynchronous, with stimuli presented at one of six SOAs (±300, ±200, and ±100 ms). Visual stimuli were presented on a BENQ XL2720-B monitor at 100 Hz, using an NVIDIA GeForce GTX 1060 (6GB) graphics card. Auditory stimuli were presented via headphones, using the PsychPortAudio functions in MATLAB.

On the first trial, participants were asked to report which of the two stimuli had occurred first using the left and right mouse buttons. On the second trial, participants were asked to report whether the Gabor patch was oriented clockwise or counter-clockwise relative to vertical. These tasks alternated throughout the experimental blocks (i.e., participants never made two successive judgements of the same type). A text prompt appeared on-screen to remind participants of the response-mapping for each trial. Response mappings were constant for both judgements throughout the experiment, and across participants. We took this approach to avoid making the task too difficult for participants, and to make analysis of the bias itself simpler, though other studies have used random response mappings between trials (e.g., Pape et al., 2017).

Each experimental block (241 trials each) took approximately 15 minutes to complete, and participants took a short break between blocks. We constructed a trial order for each block, and for each participant, which approximated orthogonal sampling of SOAs in TOJ trials relative to both orientation and SOA in orientation judgement trials. As a result, physical temporal order was approximately random with respect to physical orientation. With three blocks of 240 analysed trials (the first trial from each block was excluded from analyses), we yielded 60 TOJs for each SOA, and 60 orientation judgements for each angle. When split by preceding-SOA, we yielded ~30 TOJs per SOA in each of the preceding-SOA distributions.

Participants were given written and verbal task instructions, and completed a demonstration version of the task. This demo version of the task was identical to the experimental version, except that it was shorter (25 trials), and participants were given explicit feedback after each trial. We included this demo version as we reasoned that interleaving two psychophysical tasks here would be even more difficult than the single task used in Experiment One. Despite this demo block, 10 participants were removed from analyses as their responses indicated extremely poor performance, plausibly due to misunderstanding the task instructions or difficulty alternating between judgements.

Since the custom stimulus-presentation device used in Experiment One was limited to flashes and beeps, we used Psychtoolbox in MATLAB to conduct this experiment. One major concern using Psychtoolbox for timing experiments is the precision and accuracy of when stimuli are presented. On frames where the visual stimuli were drawn, we included a small white disc in the bottom left corner of the screen. When this frame was presented, the disc illuminated the lens of a photodiode attached to the corner of the computer monitor, providing a reliable timestamp for the onset of the visual stimulus. The audio-output from the testing PC was split, with one connection going to headphones worn by the participant, and one to an independent microcontroller running custom software. The output of the photodiode and PC-audio were recorded by the microcontroller, which determined when the stimuli had onset, returning time-stamps to the PC via USB. Stimulus timing was accurate to within ±8 ms, and no trials were excluded on these grounds.

### Data Analysis

The analytic method used here was identical to that used in Experiment One. We split participants’ TOJs into two datasets based on the temporal order of stimuli presented in the preceding trial. Of course, the preceding trial in this experiment was actually an orientation judgement, and participants were aware that they could ignore the auditory stimulus. We computed participants’ PSS and JND for both datasets, and the difference in their PSS’ to compute rapid recalibration. To quantify any response bias, we also completed this process with TOJs split by the preceding orientation judgement and preceding TOJ.

## Experiment Two results

Before considering rapid recalibration or response biases, we first computed participants’ sensitivity and accuracy to temporal order and orientation. Unfortunately, a substantial proportion of the sample had JNDs exceeding the sampling range in the temporal order judgement task, indicating they could not reliably discern the temporal order of even the most asynchronous stimuli (offset by 300 ms). Of the 30 participants recruited, 20 were included in the final analyses (one withdrew before completing the experiment, and nine were excluded for having JNDs > 300 ms). Of those remaining, there was a slight bias to perceive synchrony for audio-leads (mean PSS = −53.15 ms, SD = 84.45 ms), and similar JNDs to Experiment One (mean JND = 143.86 ms, SD = 47.28 ms). Response distributions, collated over those participants whose data were analysed, are presented in Figure 8 (below).

**Figure 8.**
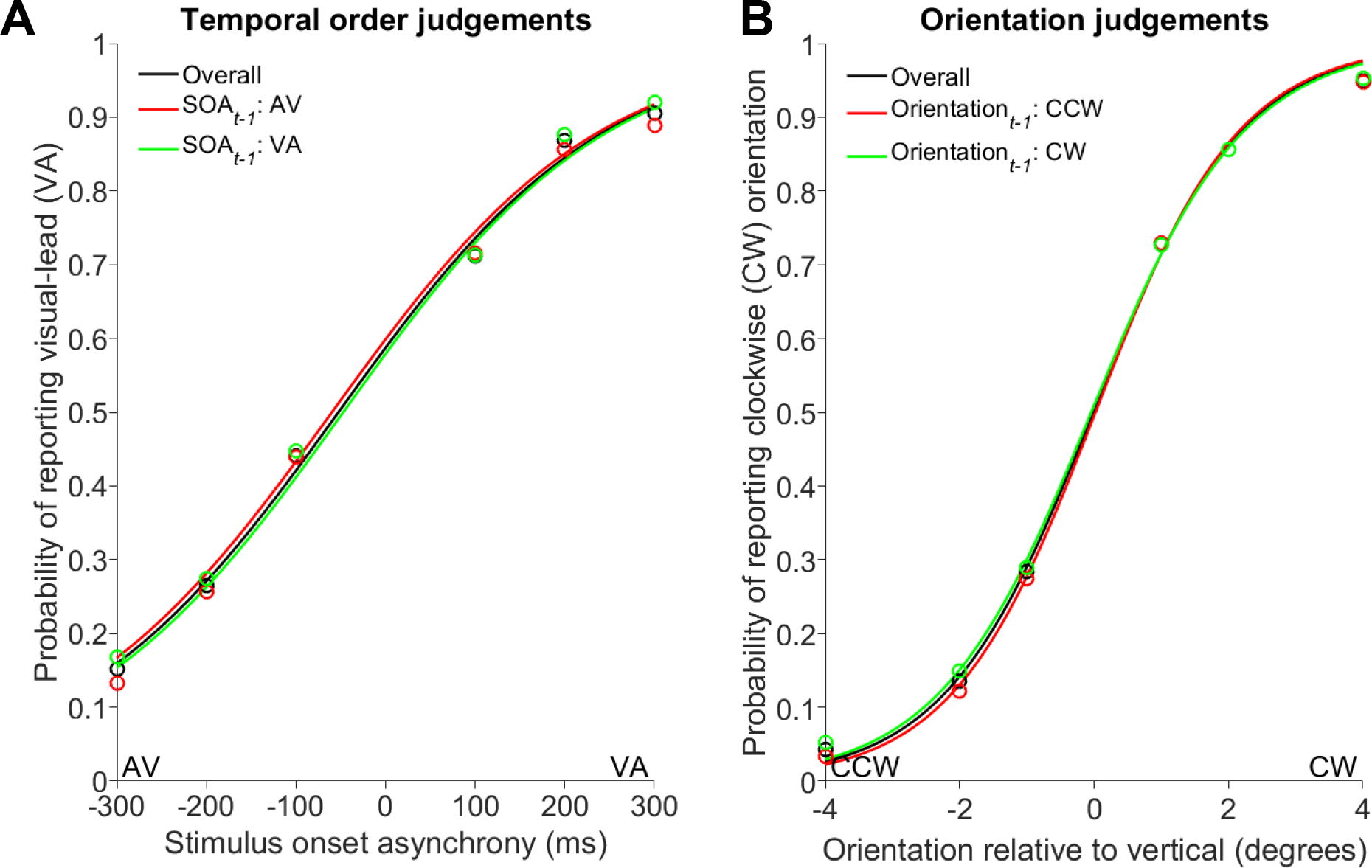
*Temporal order and orientation judgements, collated across participants*. Plotting conventions are the same as those used in Figure 1. **(A)** Participants’ TOJs were generally sensitive to the timing of stimuli. Unlike in Experiment One, we found evidence that temporal order judgements *positively* rapidly recalibrated. Note that the distributions provided here do not illustrate the within-participant differences we were testing, but provide the clearest depiction of the overall response distributions, and mimic the figure style of Van der Burg et al. (2013). **(B)** Participants’ orientation judgements, both overall and as functions of the preceding orientation. We found no evidence of a trial-order effect on orientation perception (perhaps surprisingly given the results of Fischer and Whitney, 2014), though this is beyond the scope of this paper.

Paired *t*-tests indicated that the temporal order of stimuli on the preceding trial systematically changed participants’ perception of synchrony (*t*_19_ = 2.34, *p* = .030), consistent with the effect found in SJ tasks (Simon et al., 2017; Van der Burg et al., 2013, 2015). Participants’ overall PSS was −53.15 ms; following AV trial, the average PSS was −62.51 ms, and −44.40 ms following VA trials. The magnitude of this effect (~18 ms) is consistent with that found by Van der Burg et al. (2013) in their SJ task (~20 ms). As predicted, stimulus order on the preceding trial did not change participants’ sensitivity to temporal order (*t*_19_ = 0.38, *p* = .708). Participants’ sensitivity to temporal order was uncorrelated with the magnitude of their rapid recalibration effect (*r_18_* = 0.41, *p* = .070), although this approached significance.

As a manipulation check, we then split participants’ TOJs based on their preceding orientation judgement (*trial−1*) and TOJ (*trial−2*). Paired *t*-tests revealed no evidence that participants’ TOJs were biased by their most recent button-press (i.e., orientation judgement; *t_19_* = 1.11, *p* = .279), replicating the result of this analysis in Experiment One, where TOJs were not biased by their preceding confidence judgement.

Similarly, we found no evidence that participants’ TOJs were biased by their most recent TOJ (*t_19_* = 0.88, *p* = .388). This indicates our paradigm successfully disrupted the confounding correlation between the previous TOJ and the previous SOA in Experiment One and Roseboom (2019).

## General discussion

In this paper we aimed to resolve the key issue of whether *negative* rapid recalibration of temporal order perception reflects a choice-repetition bias, unrelated to timing perception. Previous studies found that participants’ timing judgements are biased by the stimulus-order in the preceding trial. In the SJ task, participants tend to perceive repeated stimulus-order in consecutive trials as increasingly synchronous (e.g., Van der Burg et al., 2013; Simon et al., 2017). In the TOJ task (and magnitude judgement task, which requires temporal order be reported via an SOA-estimate), the opposite result was reported (Roseboom, 2019). Further analysis of Roseboom’s data (see Figure 3) indicated a bias to repeat consecutive TOJs could account for the reported effect. In essence, the previous SOA and previous TOJ were confounded.

In our first experiment we aimed to determine whether timing perception had been rapidly recalibrated, or if trial-by-trial changes in TOJs were due to a choice-repetition bias. We found no evidence that TOJs positively *or* negatively rapidly recalibrated. However, we found that participants’ confidence *positively* rapidly recalibrated. This suggests that temporal order perception had *positively* rapidly recalibrated, meaning similar SOAs in consecutive trials seemed more synchronous (or less ordered, and therefore judged with lowed confidence). We proposed that this effect would have been detectable in TOJs had it not been obfuscated by a larger, *negative* choice-repetition bias.

Experiment Two tested this prediction by dissociating the previous TOJ from the previous SOA. In this case, participants’ responses on the previous trial were independent of the SOA presented on that trial, statistically eliminating the effect of the choice-repetition bias. Under these conditions, we found evidence of *positive* rapid recalibration of TOJs, consistent with the effect reported in SJ tasks. That is, participants perceived similar SOAs in consecutive trials as more synchronous.

This result potentially undermines some of the conclusions drawn by Roseboom (2019) to explain the contradictory effects in TOJ and SJ tasks. Based on his findings, he argued that rapid recalibration likely reflects changes in decision-making processes, and these changes manifest differently for synchrony and temporal order/magnitude judgements. This was a reasonable interpretation for the set of contradictory results, and may still be true. However, our findings suggest that more traditional explanations, which assume that rapid recalibration occurs in the sensory domain, are once again plausible.

If we assume that timing information is represented by some sensory process that then feeds forward into separate decision-making processes, the most parsimonious explanation for a common effect in all judgements is that the effect is on sensory processing itself. For instance, single-trial adaptation of a population of delay-tuned neurons, like that proposed in Roach et al. (2011), could account for this effect. Of course, this is at odds with the results of Simon et al. (2017) who investigated rapid recalibration of SJs using EEG. They found that rapid recalibration was indexed by late-evoked, and therefore presumably post-sensory, processes. The alternative, then, is that a common rapid recalibration effect across timing judgements reflects a common decisional strategy.

For instance, Roseboom (2019) proposed that an assimilative effect on decision criteria could produce rapid recalibration as observed in the SJ task, in line with a proposal by Yarrow et al. (2011). He then pointed out how this would not account for his *negative* TOJ rapid recalibration effects, as such a model can produce only *positive* rapid recalibration effects. However, since we found *positive* rapid recalibration of TOJs, we should reconsider this assimilative model. A shift of decision criteria toward the SOA of recent stimulus-pairs could account for the similar effects of rapid recalibration on temporal order and synchrony judgements. This is also consistent with the observed changes in participants’ confidence reports.

To clarify the point, imagine a participant perceives a flash-lead SOA on some trial, and their decision criterion then shifts slightly toward flash-lead SOAs. If the subsequent trial is a flash-lead SOA, it is less likely to be perceived as such, being closer to the criterion, accounting for the effect observed here. If we assume that confidence is in some way related to the difference in participants’ perceived SOA and the decision criterion, the second trial would also be associated with lower confidence, as observed in Experiment One. This simple decisional model, relying on assimilation of decision criteria, provides neat and supported predictions regarding trial-by-trial changes in temporal order and confidence judgements. However, it is not sufficient to explain other recent findings (e.g., Roseboom et al., 2015), making it unlikely that changes in decisional processing solely drive rapid recalibration (or temporal recalibration following prolonged adaptation).

Roseboom et al. (2015) found that prolonged exposure to temporal asynchrony induces changes not only in synchrony perception, but the sensitivity of the observer to timing information itself. For instance, when participants are repeatedly exposed to a particular timing relationship, their sensitivity to other timing relationships is relatively increased. This effect is at odds with simple, purely decisional, accounts of temporal recalibration. It remains to be seen whether such effects (non-uniform distortion of sensitivity around adapted SOAs) are observable on the scale of a single trial. If so, that would provide strong evidence that rapid recalibration occurs at the level of sensory processing.

## Conclusion

This study aimed to determine whether rapid recalibration of temporal order perception reflects a generalised bias in decision making, unrelated to timing perception. Analyses of our own data and data provided by Roseboom (2019) indicated that a simple bias to repeat judgements in consecutive trials could account for the contradictory effects reported in temporal order and synchrony judgement tasks. In our first experiment, we measured participants’ confidence in their order judgements and found evidence of *positive* rapid recalibration. In a final experiment that employed a novel design with interleaved tasks, we eliminated participants’ bias to repeat their judgements and found evidence that temporal order judgements *positively* rapidly recalibrate, consistent with synchrony judgements. It remains unclear how rapid recalibration manifests, and at what level of processing; this effect could be accounted for by simple changes in either sensory or decision-making processes. Further research is required to determine the locus of these effects, though attempting to replicate recent findings by Roseboom et al. (2015) in single-trial adaptation would appear to be a good place to start.

## References

Abrahamyan, A., Silva, L. L., Dakin, S. C., Carandini, M., & Gardner, J. L. (2016). Adaptable history biases in human perceptual decisions. Proceedings of the National Academy of Sciences, 113(25), E3548–E3557.

Binder, M. (2015). Neural correlates of audiovisual temporal processing-comparison of temporal order and simultaneity judgments. Neuroscience, 300, 432–447.

Cross, D. V. (1973). Sequential dependencies and regression in psychophysical judgments. Perception & Psychophysics, 14(3), 547–552.

Donohue, S. E., Woldorff, M. G., & Mitroff, S. R. (2010). Video game players show more precise multisensory temporal processing abilities. Attention, Perception, & Psychophysics, 72(4), 1120– 1129.

Fischer, J., & Whitney, D. (2014). Serial dependence in visual perception. Nature Neuroscience, 17(5), 738.

Fründ, I., Wichmann, F. A., & Macke, J. H. (2014). Quantifying the effect of intertrial dependence on perceptual decisions. Journal of Vision, 14(7), 9–9.

Fujisaki, W., Shimojo, S., Kashino, M., & Nishida, S. Y. (2004). Recalibration of audiovisual simultaneity. Nature Neuroscience, 7(7), 773.

García-Pérez, M. A., & Alcalá-Quintana, R. (2012). On the discrepant results in synchrony judgment and temporal-order judgment tasks: a quantitative model. Psychonomic Bulletin & Review, 19(5), 820–846.

Hwang, E. J., Dahlen, J. E., Mukundan, M., & Komiyama, T. (2017). History-based action selection bias in posterior parietal cortex. Nature Communications, 8(1), 1242.

Keane, B., Spence, M., Yarrow, K., & Arnold, D. (2015). Perceptual confidence demonstrates trial-by-trial insight into the precision of audio–visual timing encoding. Consciousness and Cognition, 38, 107–117.

Kiyonaga, A., Scimeca, J. M., Bliss, D. P., & Whitney, D. (2017). Serial dependence across perception, attention, and memory. Trends in Cognitive Sciences, 21(7), 493–497.

Kondo, A., Takahashi, K., & Watanabe, K. (2012). Sequential effects in face-attractiveness judgment. Perception, 41(1), 43–49.

Love, S. A., Petrini, K., Cheng, A., & Pollick, F. E. (2013). A psychophysical investigation of differences between synchrony and temporal order judgments. PloS one, 8(1), e54798.

Marinovic, W., Poh, E., de Rugy, A., & Carroll, T. J. (2017). Action history influences subsequent movement via two distinct processes. Elife, 6, e26713.

Roach, N. W., Heron, J., Whitaker, D., & McGraw, P. V. (2010). Asynchrony adaptation reveals neural population code for audio-visual timing. Proceedings of the Royal Society B: Biological Sciences, 278(1710), 1314–1322.

Roseboom, W. (2019). Serial dependence in timing perception. Journal of Experimental Psychology: Human Perception and Performance, 45(1), 100.

Roseboom, W., Linares, D., & Nishida, S. Y. (2015). Sensory adaptation for timing perception. Proceedings of the Royal Society B: Biological Sciences, 282(1805), 20142833.

Simon, D. M., Noel, J. P., & Wallace, M. T. (2017). Event related potentials index rapid recalibration to audiovisual temporal asynchrony. Frontiers in Integrative Neuroscience, 11, 8.

Stekelenburg, J. J., Sugano, Y., & Vroomen, J. (2011). Neural correlates of motor-sensory temporal recalibration. Brain Research, 1397, 46–54.

Van der Burg, E., Alais, D., & Cass, J. (2013). Rapid recalibration to audiovisual asynchrony. Journal of Neuroscience, 33(37), 14633–14637.

Van der Burg, E., Orchard-Mills, E., & Alais, D. (2015). Rapid temporal recalibration is unique to audiovisual stimuli. Experimental Brain Research, 233(1), 53–59.

Van Eijk, R. L., Kohlrausch, A., Juola, J. F., & van de Par, S. (2008). Audiovisual synchrony and temporal order judgments: Effects of experimental method and stimulus type. Perception & Psychophysics, 70(6), 955–968.

Verstynen, T., & Sabes, P. N. (2011). How each movement changes the next: an experimental and theoretical study of fast adaptive priors in reaching. Journal of Neuroscience, 31(27), 10050–10059.

Vroomen, J., Keetels, M., De Gelder, B., & Bertelson, P. (2004). Recalibration of temporal order perception by exposure to audio-visual asynchrony. Cognitive Brain Research, 22(1), 32–35.

Yarrow, K., Jahn, N., Durant, S., & Arnold, D. H. (2011). Shifts of criteria or neural timing? The assumptions underlying timing perception studies. Consciousness and Cognition, 20(4), 1518–1531.

